# Characterisation of Peripheral Blood B Cell Receptor Repertoire in Severe Eosinophilic Asthma and EGPA

**DOI:** 10.64898/2026.06.16.732558

**Authors:** Jantarika Kumar Arora, Emily Bessell, Sonay Beyatli, Delphine Thenet, James Brown, Ahuva Nissim, Myles J. Lewis, Louisa K. James, Paul E. Pfeffer

## Abstract

**Background:** Severe eosinophilic asthma (SEA), eosinophilic granulomatosis with polyangiitis (EGPA) and nasal polyposis (NP) are immune-mediated diseases characterised by eosinophilic inflammation. However, there is also increasing interest in the potential pathological roles of autoantibodies in these diseases. Understanding their B cell receptor (BCR) repertoires may provide valuable insights into disease mechanisms, and potential role of B cells in their pathology.

**Methods:** We conducted BCR repertoire sequencing using peripheral blood from 43 patients, comprising SEA with nasal polyps (SEA+NP), SEA without nasal polyps (SEA-NP), and EGPA, along with 16 healthy controls (HCs).

**Results:** Compared to HCs, patients with EGPA exhibited increased relative proportions of IgA1, IgG1, IgG2, and IgG4 subclasses. Similarly, SEA-NP patients demonstrated significantly high proportion of IgG2 sequences. Notably, the IgG4 subclass was significantly elevated across all patient groups compared to HCs. Patients receiving anti-IL-5/5R biologic treatments showed increased relative proportions of IgA2 and IgG2 subclasses compared to untreated patients. Some variation across participant groups in mean somatic hypermutation and mutation frequency was evident. 1,508 clones shared across patients, but not healthy controls, were evident though the majority showed low clonal expansion. Nevertheless, a few shared clones did show either high prevalence across patients and/or higher clonal expansion.

**Conclusion:** Changes in BCR repertoires in SEA/EGPA are consistent with a pattern of a more mature B cell component in the periphery and with the T2 inflammatory response observed in SEA and EGPA. BCR clonotypes shared across patients were evident, however, whether such clonotypes are pathological in SEA/EGPA requires further investigation.

## Background

Aberrant T2-type inflammation is central to the pathology of severe eosinophilic asthma (SEA),[1] yet the underlying cause of persistent inflammation remains unclear. The proportion of patients with underlying atopy decreases with increasing age and severity, suggesting that in most patients continuing exposure to an environmental allergen is not a driving cause of severe asthma.[2] In the closely-related pathology of eosinophilic granulomatosis with polyangiitis, EGPA, approximately 30–40% of patients have evidence of clinically-relevant anti-neutrophil cytoplasmic antibodies (ANCA), typically anti-myeloperoxidase (anti-MPO) autoantibodies.[3] [4] Research over recent years has described an increasing number of auto-antibodies present at increased concentration in serum,[5] and anti-eosinophil peroxidase (EPX) antibodies in sputum, in patients with severe asthma.[6] It remains uncertain whether such autoantibodies are directly pathological or an epiphenomenon. Whilst anti-IL-5 monoclonal antibody biologic therapies are effective in treating SEA and EGPA,[7] [8] anti-B-cell biologic therapy with Rituximab is effective in many patients with EGPA,[9] suggesting that B cells have a major role in that pathology though their role in severe eosinophilic asthma is less clear.

We previously investigated seroprevalence of autoantibodies in patients with SEA and EGPA,[5] and we now describe the B cell receptor (BCR) repertoire of peripheral blood B cells in these patients. BCR gene transcripts encode the antibodies that individual B cells will produce, both the constant region encoding immunoglobulin isotype and the variable region responsible for antigen binding specificity. Importantly, different immunoglobulin isotype have significant biologic differences [10]. Furthermore, clonality of BCR sequences (clonotypes) present in patients with disease, but not healthy controls, can suggest disease-related antibodies such as might be apparent in autoimmune disease or in response to infection [11] [12] [13].

We aimed to characterize whether the repertoire properties (isotype distribution, somatic hypermutation, and clonal expansion) differed in patients with SEA/EGPA compared to healthy controls and whether specific and distinct clonotypes were present in peripheral blood in patients with SEA/EGPA. Better understanding of the B cell repertoire in severe asthma and EGPA will help us understand their pathogenesis and potentially yield diagnostic/ theragnostic biomarkers.

## Methods

### Patient Sample Collection

Samples of peripheral blood mononuclear cells (PBMCs) from 60 participants, from the prior phase of the study [5], were selected by purposive sampling to ensure a broad range of treatable trait characteristics. The sample from one individual did not pass quality assurance and was therefore excluded from analysis – samples from 59 participants were analysed. Participants for these analyses were of four types – healthy controls (HCs), patients with severe eosinophilic asthma with nasal polyps (SEA+NP), patients with severe eosinophilic asthma without nasal polyps (SEA-NP), and patients with eosinophilic granulomatous polyangiitis (EGPA). Further details about patient recruitment and characterization are as previously described [5]. Recruitment and analysis were conducted after ethics committee approval (NHS REC ethics approval 20/PR/0004).

### Bulk BCR repertoire library preparation and sequencing

Total RNA was extracted using the RNeasy Mini RNA Isolation Kit (Qiagen) according to the manufacturer’s protocol. Heavy chain variable region sequences (*IGHV*) were amplified using isotype-specific primers containing unique molecular identifiers (UMIs) of 10 or 12 nucleotides as previously described [12]. Briefly, 200 to 500 ng of RNA was annealed at 72°C for 5 minutes, followed by incubation on ice for 2 minutes. First-strand complementary DNA (cDNA) was synthesized using SuperScript IV Reverse Transcriptase (Thermo Fisher Scientific). Second-strand cDNA synthesis was performed using Phusion High-Fidelity (HF) DNA Polymerase (New England Biolabs) and a pool of UMI-containing IGHV forward primers (98°C for 4 minutes, 52 °C for 1 minute, and 72°C for 5 minutes). Double-stranded cDNA was purified twice using Ampure XP beads (Beckman Coulter) according to manufacturer’s instructions before library amplification with Platinum Taq HF Polymerase (Life Technologies). Libraries were quantified using the Qubit dsDNA High Sensitivity Assay Kit and Bioanalyzer 2100 Expert High Sensitivity DNA Assay before Illumina MiSeq sequencing with paired-end 301 base pair (bp) reads.

### Bulk BCR repertoire processing, quality assurance and analysis

Raw sequencing data were assessed for overall sequencing qualities using FastQC (V0.11.9) (https://www.bioinformatics.babraham.ac.uk/projects/fastqc/). Unique molecular identifier (UMI)-collapsed consensus VDJ sequences were generated and processed using pRESTO (v0.7.2) (https://pmc.ncbi.nlm.nih.gov/articles/PMC4071206/). Paired-end reads with mean Phred quality scores below 20 were excluded. UMI barcodes were aligned using MUSCLE (v3.8.31) (https://academic.oup.com/nar/article/32/5/1792/2380623), and UMIs with more than three unique reads were used to generate consensus VDJ sequences. Paired-end UMI consensus sequences were then assembled into a single VDJ contig, and constant region isotype were annotated using MaskPrimers.py in pRESTO (v0.7.2). Duplicate VDJ sequences were collapsed using CollapseSeq.py in pRESTO (v0.7.2). VDJ genes were annotated using AssignGenes.py (ChangeO v1.3.0) (https://academic.oup.com/bioinformatics/article/31/20/3356/195677) and IgBLAST (v1.22) (https://academic.oup.com/nar/article/41/W1/W34/1097536), followed by correction of V alleles using TIgGER (v.1.1.0) (https://www.pnas.org/doi/10.1073/pnas.1417683112). Clonally related sequences, defined as those sharing the same V gene, J gene and CDR3 junction length, were identified using scoper (v1.3.0) (https://doi.org/10.1371/journal.pcbi.1007977). Germline sequences were inferred using CreateGermlines.py, followed by calculation of somatic hypermutation using observedMutations function in shazam (v1.2.0). Clonal diversity analysis was performed using the alphaDiversity function from alakazam (v1.3.0). Diversity was calculated over a range of diversity orders from 0 (min_q=0) to 4 (max_q=4) in 0.1 increments of 0.1 (step_q=0.1), with 95% confidence intervals (ci=0.95). A total of 100 bootstrap resampling realizations (nboot=100) were applied to generate a smooth curve.

### Convergent and phylogenetic tree analyses

Clonal relationships of amino acid sequences in each group were identified via hierarchical clustering in scoper (v1.3.0), with a mean threshold of 0.06762372 calculated via a smoothed density approach implemented in shazam (v1.2.0). Clones with a size greater than one and present in at least two samples per group were combined before defining cross-group clones using the identicalClones function in scoper (based on identical V gene, J gene and CDR3 junction length). Convergent clones were defined as those present in patient groups but absent in healthy controls. The lineage trees of convergent clones were reconstructed via maximum parsimony using buildPhylipLineage from alakazam (v1.3.0).

### Statistical analysis

Differences in the percentage of unique isotypes and mean somatic hypermutation frequencies were compared using the Kruskal-Wallis test. For pairwise comparisons between two independent groups, the Wilcoxon rank-sum test (Mann-Whitney) was applied. The number of samples for each analysis is indicated in the figure legends. The significance levels are defined as follows: ns = p > 0.05, ∗p ≤ 0.05, ∗∗p ≤ 0.01, ∗∗∗p ≤ 0.001, and ∗∗∗∗ p ≤ 0.0001.

## Results

Peripheral blood BCR repertoires were bulk sequenced from 16 healthy controls (HCs, mean age 42.2 yrs, 50% female), 15 SEA–NP (mean age 52.9 yrs, 47% female), 16 SEA+NP (mean age 56.1 yrs, 44% female) and 12 EGPA donors (mean age 56.3 yrs, 50% female) (Table 1).

**Table 1:** Participant characteristics.

|  | Healthy Control | SEA-NP | SEA+NP | EGPA |
| --- | --- | --- | --- | --- |
| Number of Participants | 16 | 15 | 16 | 12 |
| Gender |  |  |  |  |
| Female | 8 | 7 | 7 | 6 |
| Male | 8 | 8 | 9 | 6 |
| Mean age /yrs | 42.2 | 52.9 | 56.1 | 56.3 |
| Smoking status |  |  |  |  |
| Never Smoker | 14 | 9 | 7 | 7 |
| Ex-smoker | 1 | 6 | 9 | 3 |
| Current smoker | 0 | 0 | 0 | 0 |
| Current Treatment |  |  |  |  |
| anti-IL-5/5R | 0 | 4 | 7 | 3 |
| systemic corticosteroids | 0 | 5 | 1 | 4 |

### Relative distribution of antibody subclasses

The relative distribution of antibody subclasses varied across patient groups (Figure 1A). IgA1 and IgG1 showed significantly higher proportions in patients with EGPA compared to HCs, whereas IgG2 was elevated in both SEA–NP and EGPA (Figure 1A). IgE was more abundant across all three patient groups with statistical significance observed with SEA+NP, while IgG4 showed a significant increase across all patient groups compared to HCs (Figure 1A). Conversely, the relative proportion of IgM was lower across all patient groups compared to HCs (Figure 1A).

**Figure 1.**
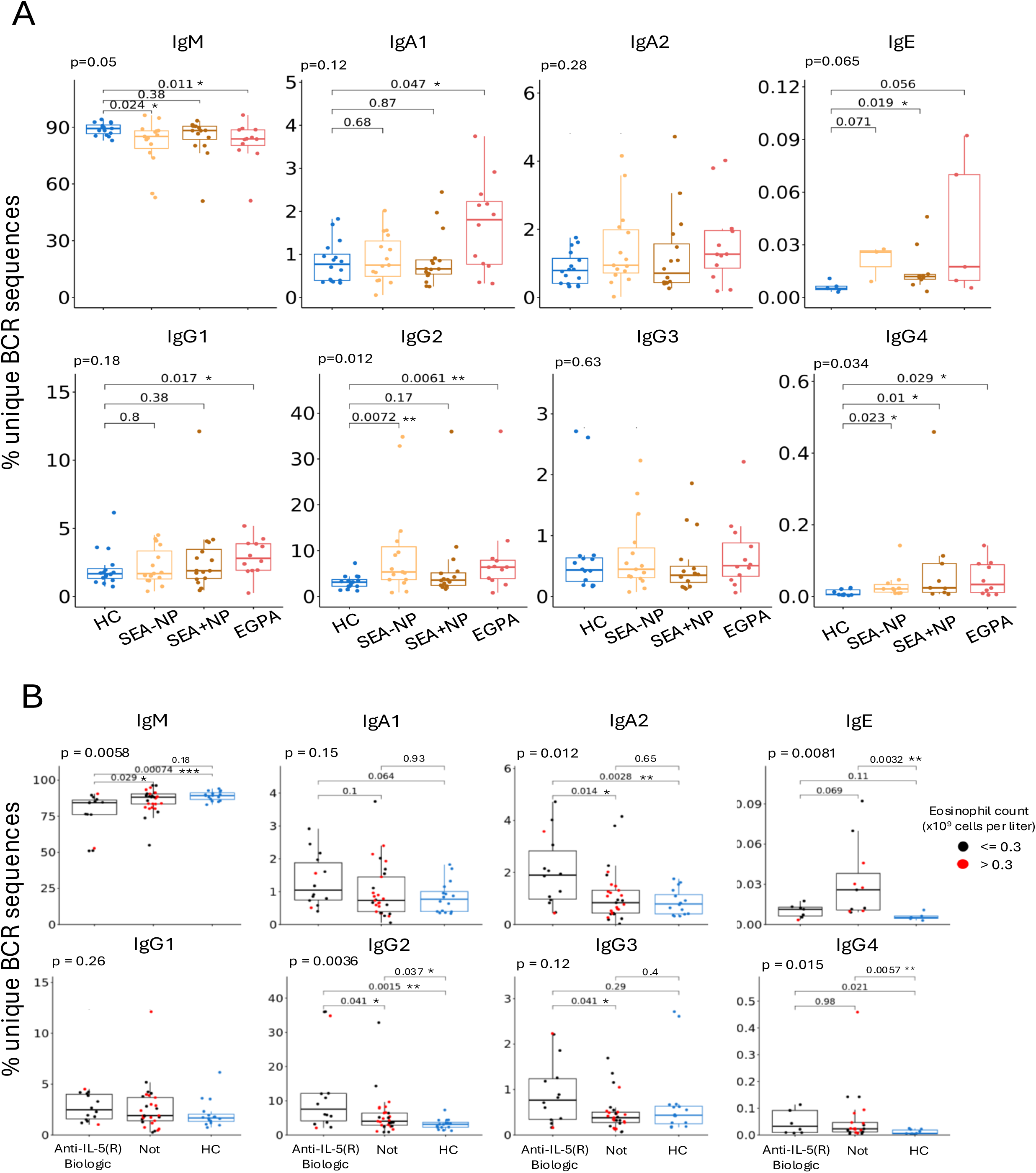
Relative distribution of immunoglobulin subclass characteristics in HC, SEA–NP, SEA+NP, and EGPA participants. **A,** Mean frequency of immunoglobulin subclasses compared between patients with different disease phenotypes and healthy controls. **B,** compared between patients on and not on anti-IL-5/5R biologics. Significance levels are defined as follows: ns = p > 0.05, ∗p ≤ 0.05, ∗∗p ≤ 0.01, ∗∗∗p ≤ 0.001, and ∗∗∗∗ p ≤ 0.0001. *HC,* Healthy Control; *SEA –NP,* Severe eosinophilic asthma without nasal polyps; *SEA +NP,* Severe eosinophilic asthma with nasal polyps; *EGPA,* Eosinophilic granulomatosis polyangiitis.

Comparing those patients on anti-IL-5/5R biologics to those not, significantly increased frequencies of IgA2, IgG2 and IgG3 were evident in patients on anti-IL-5/5R biologics (Figure 1B). A significantly decreased frequency of unique IgM sequences and trend towards decreased IgE sequences was also evident in patients on anti-IL-5/5R biologics (Figure 1B). In patients not on anti-IL-5/5R biologics, there was no clear association between (sub)class frequencies and the most recent peripheral blood eosinophil count.

### Differences in clonal diversity

Significant differences in mean somatic hypermutation (SHM) frequency, calculated from individual clones, were evident for IgA2, IgG1 and IgG3 (Figure 2A). Clones in IgA2 were less mutated in patient groups compared to HCs, whereas IgG1 and IgG3 clones exhibited higher SHM frequencies across patient groups, with statistical significance observed in SEA-NP compared to HCs (Figure 2A).

**Figure 2.**
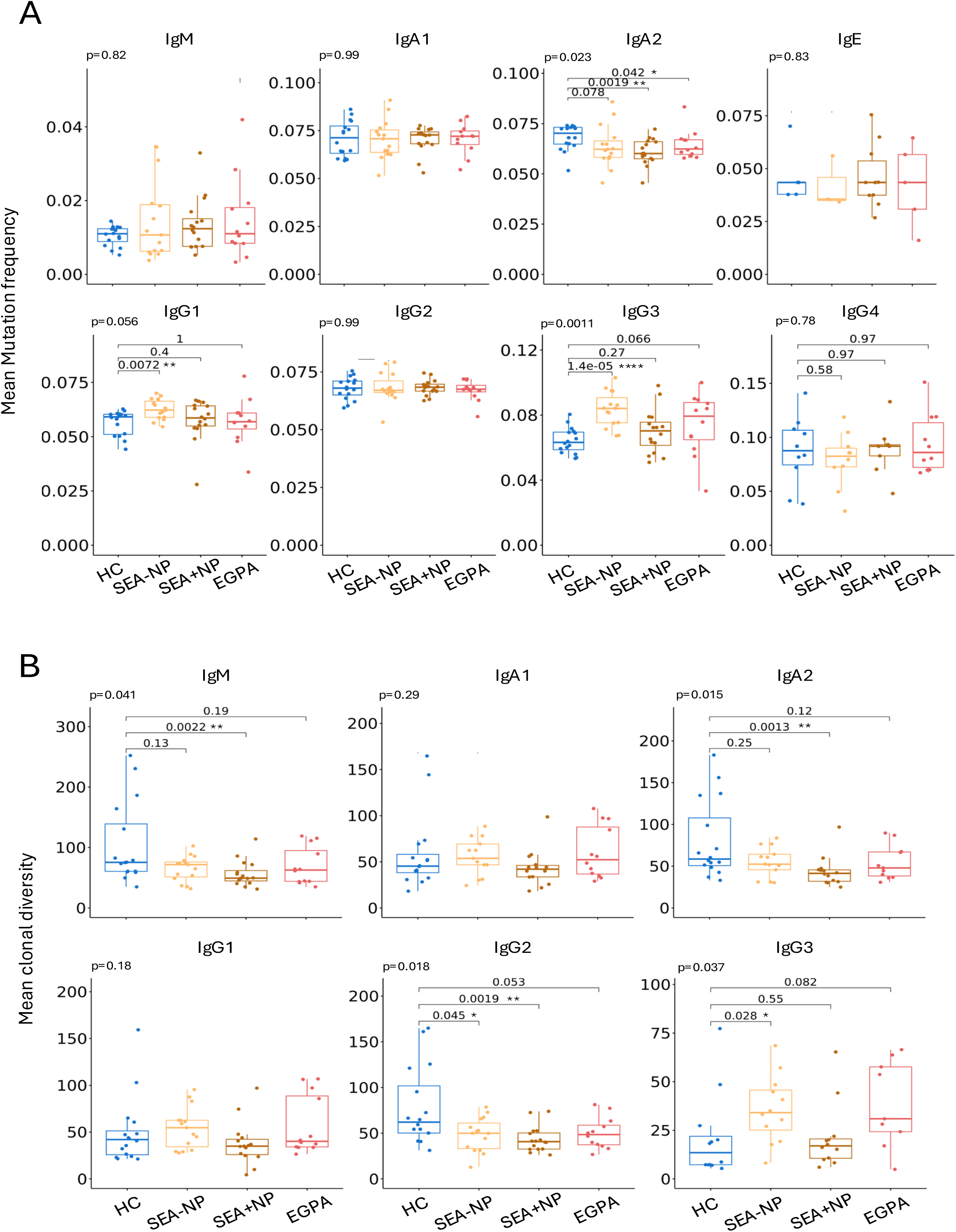
Somatic hypermutation and clonal diversity of B cell receptors in HC, SEA–NP, SEA+NP, and EGPA participants. **A,** Mean somatic hypermutation frequency per donor, calculated from individual clones. **B,** Mean clonal diversity per donor, calculated from individual clones. Significance levels are defined as follows: ns = p > 0.05, ∗p ≤ 0.05, ∗∗p ≤ 0.01, ∗∗∗p ≤ 0.001, and ∗∗∗∗ p ≤ 0.0001. *HC,* Healthy Control; *SEA –NP,* Severe eosinophilic asthma without nasal polyps; *SEA +NP,* Severe eosinophilic asthma with nasal polyps; *EGPA,* Eosinophilic granulomatosis polyangiitis.

IgM, IgA2, and IgG2 exhibited lower clonal diversity in patient groups, with statistical significance in the SEA+NP group, relative to HCs (Figure 2B), indicating greater clonal expansion within these subclasses. In contrast, IgG3 showed higher clonal diversity in patients, with a significant difference observed in the SEA–NP compared to HCs, suggesting less clonal expansion within this subclass (Figure 2B).

### Presence of shared clonotypes

We compared amino acid sequences of expanded clones across participants and found that 1,508 clones were present in more than one patient but absent in HCs (Figure 3A). Of these, 922 clones were observed in SEA-NP patients, 592 in SEA+NP patients, and 533 in EGPA patients (Figure 3A). 365 clones were shared across two patient groups, and 87 clones were detected in all three patient groups (Figure 3A). Focusing on the 87 shared clones, the number of patients sharing a given BCR clone ranged from 3 patients (one from each patient group) to 23 patients, with a median of 7 patients (Figure 3B). 28 clones were present in at least nine patients (Figure 3B).

**Figure 3.**
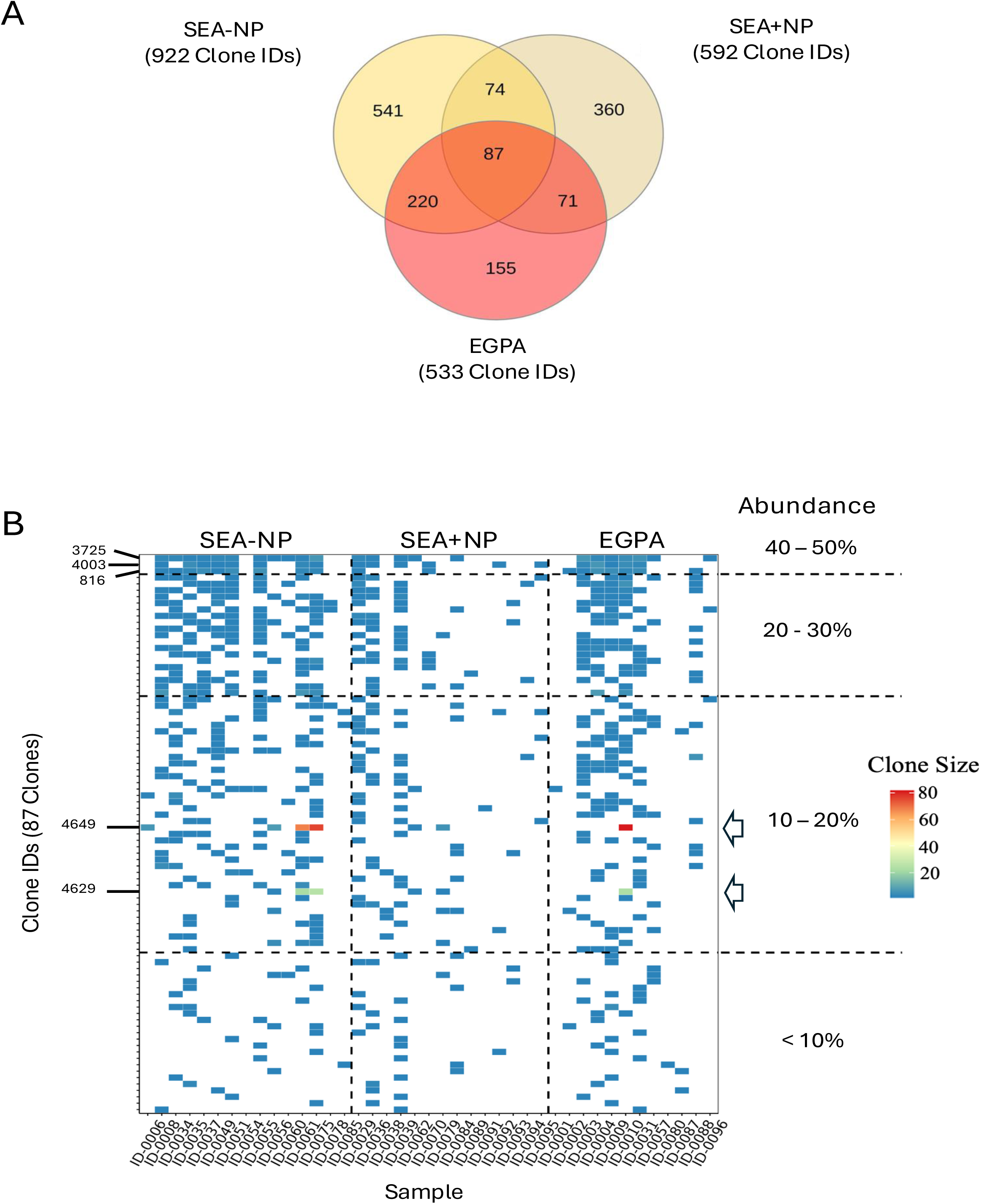
Shared B cell receptor clones patient groups at the amino acid level. **A,** Venn diagram showing overlap of B cell receptor (BCR) clones among the three patient groups. Only clones absent in HC are included. **B,** Heatmap of 87 clones shared across all three patient groups but not in HCs. Each row represents a clone, each column a donor, and color intensity indicates clone size. Percentages represent the proportion of donors sharing each clone. Open arrowheads indicating two clones showing evidence of substantial expansion. *SEA–NP,* Severe eosinophilic asthma without nasal polyps; *SEA+NP,* Severe eosinophilic asthma with nasal polyps; *EGPA,* Eosinophilic granulomatosis polyangiitis. *SHM;* somatic hypermutation.

The majority of shared clones were relatively small, containing fewer than 10 clonally related sequences in any given patient (Figure 3B), suggesting that they may not be highly expanded. Three clones (clone IDs: 3725, 4003, and 816) detected in 40 - 50% of patients but not in HCs, expressed IgM, IgG2 or IgA2 subclasses with moderate levels of SHM (Figure 3B and Figure 4A top three clones; example clone in Figure 4B). In contrast, two clones (clone IDs: 4649, and 4629) showing substantial clonal expansion were predominantly IgM with a relatively high SHM level (Figure 4A, lower 2 clones; example clone in Figure 4C).

**Figure 4.**
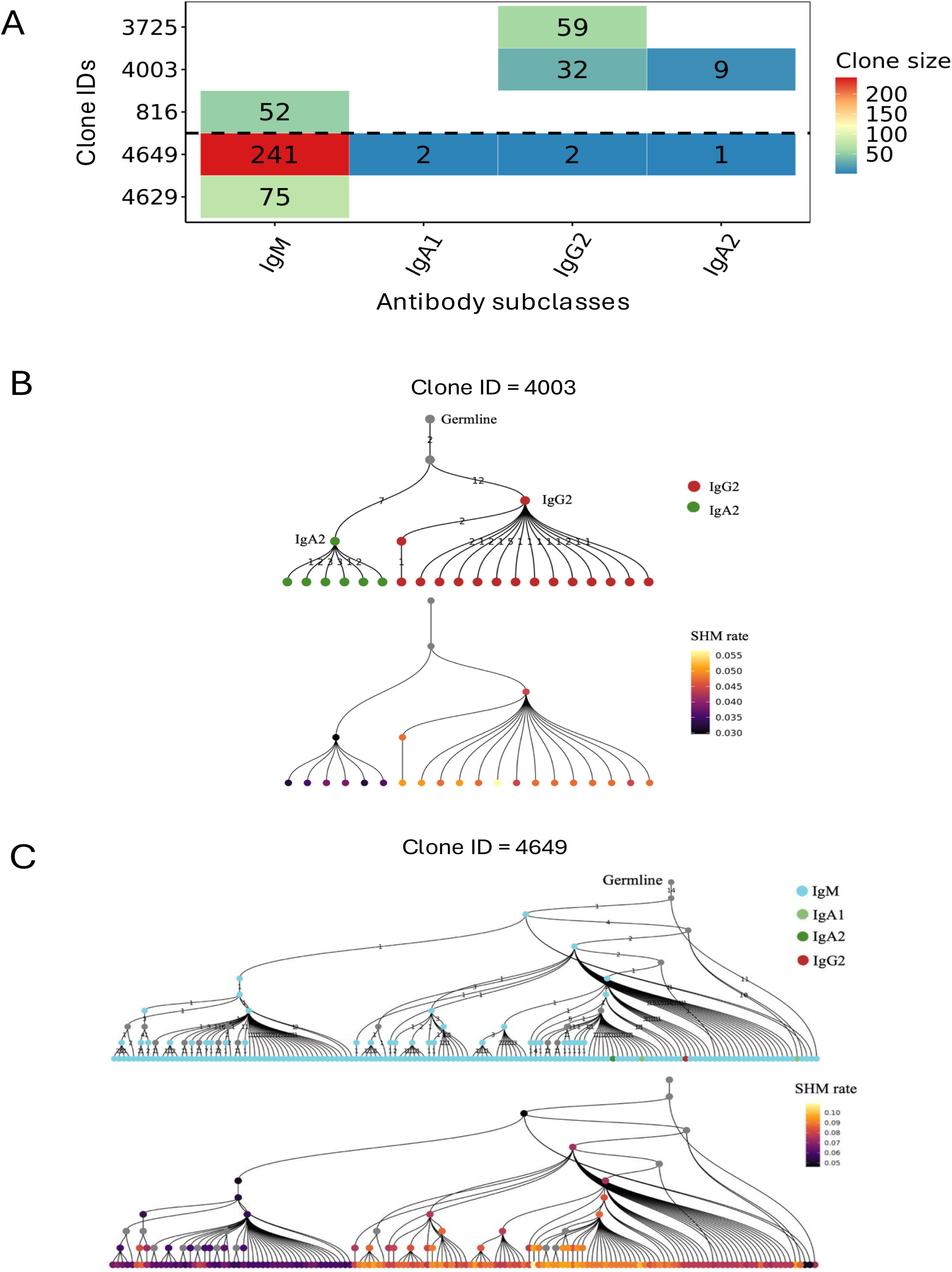
Representative B cell lineage trees in patient groups at the amino acid level. **A,** Heatmap of 5 clones illustrating distribution of BCRs across isotypes. Each row represents a clone, each column an isotype, and color intensity indicates clone size. Shows the 3 clones present in the most patients (top 3 rows) and 2 clones with substantial clonal expansion (lower 2 rows). **B,** Representative inferred B cell lineage tree for a clonotype present in 40-50% of donors (from A). **C,** Representative inferred B cell lineage tree for a clonotype present in 10-20% of donors with the highest clone size (from A). The top panel is colored by isotope; the bottom panel is colored by SHM levels. *SEA–NP,* Severe eosinophilic asthma without nasal polyps; *SEA+NP,* Severe eosinophilic asthma with nasal polyps; *EGPA,* Eosinophilic granulomatosis polyangiitis. *SHM;* somatic hypermutation.

## Discussion

Although there is increasing evidence of autoantibodies in severe asthma [5] [6] and a known association with anti-MPO autoantibodies in EGPA, [4] there has to-date been relatively little research into the role of B cells in these diseases. In order to better understand the B cell receptor (BCR) repertoire in severe asthma and EGPA, we performed BCR repertoire sequencing of peripheral blood B cells from these patients and compared characteristics of those repertoires with healthy controls. We found evidence of perturbation of peripheral BCR repertoires in patients compared to healthy controls. Firstly, in patients with severe asthma and EGPA the relative expression of antibody subclasses differed from healthy controls, with an increase in switched isotypes and a converse decrease in the relative frequency of IgM. This could indicate an increase in the proportion of memory versus naive B cells in the circulation in these patients. In support of this we also found evidence of reduced clonal diversity of some isotypes which could correspond to increased expansion of antigen-specific B cells. There was an increased frequency of somatic hypermutation in certain antibody subclasses, supporting that there may be antigen-driven clonal expansion driving this increase in circulating memory B cells in these patients. As it is not possible to determine antigen-specificity from BCR sequences, we next examined clonally-related sequences for evidence of clonal convergence across patient groups. Clonal convergence is frequently reported in response to acute infection but has also been described in patients with autoimmune disease. [11] [12] [13]

Comparing clones at the amino acid sequence level, we identified over 1500 clones that were common to patients but not healthy controls which ranged in abundance from clones present in less than 10% of patients to clones that were highly abundant and present in over 50% of the patients. The majority of identified clones shared between patients, but not shared with healthy controls, were of small clone size suggesting they may not be highly expanded. Clones that were more abundant were typically highly mutated and were restricted to the IgM, IgA2 and IgG2 subclasses potentially suggestive of bacterial-driven immune responses [10]. Future work to isolate B cells expressing these shared clonal sequences could provide more insight into the antigen/s that drive these responses which could in turn determine whether they represent a unique signature in patients with asthma/EGPA.

Few other studies have examined BCR repertoires in severe eosinophilic asthma and EGPA. Bashford-Rogers and colleagues have examined BCR repertoires in EGPA though their patient cohort was potentially characterized more by a vasculitic disease expression than a severe asthma disease expression [14]. Similar to us they found increased percentages of unique sequences of IgE and IgG4 subclasses, but in contrast also an increase in unique IgG3 sequences though their analysis was unable to distinguish between IgG1 and IgG2 subclass sequences [14]. Increased IgG4 transcripts is consistent with previous findings of elevated serum levels of IgG4 in EGPA and to a less extent in severe asthma [15]. Similarly previous research has shown elevated serum IgE in some patients with severe asthma/EGPA [16] [17].

One potential limitation of the current study is that we analyzed BCR repertoire in the peripheral blood rather than in respiratory tract tissue. Ohm-Laursen and colleagues have previously investigated BCR repertoires in blood and airway tissue from patients with atopic asthma and non-atopic controls, and found differences in BCR repertoire with more BCR mutations in tissue B cells than peripheral blood B cells from matched donors [18]. Asamori and colleagues have recently undertaken BCR sequencing to investigate autoimmunity in chronic rhinosinusitis with nasal polyposis but, in contrast to our methodology, they sequenced nasal polyp tissue to identify dominant BCR clones encoding putative autoantigens [19].

Potential explanations for differences in BCR repertoires in patients include autoantigen responses, the influence on BCR repertoires of pro-inflammatory T2 cytokines, changes in the microbiome and the effects of corticosteroid treatments. Interleukin-5 (IL-5), the major cytokine promoting eosinophilic inflammation, was originally described as a B cell growth and differentiation factor, and nasal polyp plasma cells have been shown to express the IL-5 receptor alpha subunit.[20] [21] Immunoglobulin deficiencies have been reported in some patients with asthma,[22] [23] and high exposure to systemic corticosteroids, as seen in many patients with severe asthma and EGPA, is associated with secondary antibody deficiencies [24].

The airway microbiome is known to be perturbed in asthma, which would additionally be expected to influence antibody repertoires in patients [25]. Changes in IgG2 expression and clonal diversity were in particular evident in patients - IgG2 antibodies are particularly important for antibody responses to polysaccharide antigens and immune responses to encapsulated bacteria.[26] Indeed, IgG2 deficiency is associated with more frequent exacerbations and disease progression in bronchiectasis.[27] Molecular mimicry with antibodies cross-reactive to microbial and self-antigens is thought to underlie many autoantibody responses. Indeed, many of the putative autoantibodies in nasal polyposis identified by Asamori and colleagues were cross-reactive to microbial antigens though not to encapsulated bacteria.[19]

A limitation of our research is the inability to derive specificities from BCR sequence alone – our approach cannot confirm or characterise antigen-specific pathological B cell functions in this patient group. However, recombinant expression of these clones may yet allow more insight into the role of clonal antibodies in pathologic B cell responses in SEA/EGPA pathology in the future.

In summary, significant differences in BCR repertoires are evident in patients with severe eosinophilic asthma and EGPA compared to healthy controls however further research is needed to establish the pathological and clinical significance of these differences.

## Clinical trial registration

ClinicalTrials.gov, identifier, NCT04671446.

## Abbreviations used

SEA: Severe eosinophilic asthma
EGPA: Eosinophilic granulomatosis polyangiitis
ANCA: Anti-neutrophil cytoplasmic antibodies
Anti-MPO: Anti-myeloperoxidase
Anti-EPX: Anti-eosinophil peroxidase
BCR: B cell receptor
PBMCs: Peripheral blood mononuclear cells
HC: Healthy control
SEA+NP: Severe eosinophilic asthma with nasal polyps
SEA-NP: Severe eosinophilic asthma without nasal polyps

## References

1. Jackson, D.J., P. Akuthota, and F. Roufosse, Eosinophils and eosinophilic immune dysfunction in health and disease. Eur Respir Rev, 2022. 31(163).

2. Thomsen, S.F., Epidemiology and natural history of atopic diseases. Eur Clin Respir J, 2015. 2.

3. Mahr, A., et al., Eosinophilic granulomatosis with polyangiitis (Churg-Strauss): evolutions in classification, etiopathogenesis, assessment and management. Curr Opin Rheumatol, 2014. 26(1): p. 16–23.

4. Healy, B., et al., Antineutrophil cytoplasmic autoantibodies and myeloperoxidase autoantibodies in clinical expression of Churg-Strauss syndrome. J Allergy Clin Immunol, 2013. 131(2): p. 571–6.e1-6.

5. Esposito, I., et al., Identification of autoantigens and their potential post-translational modification in EGPA and severe eosinophilic asthma. Front Immunol, 2023. 14: p. 1164941.

6. Mukherjee, M., et al., Sputum autoantibodies in patients with severe eosinophilic asthma. J Allergy Clin Immunol, 2018. 141(4): p. 1269–1279.

7. Jackson, D.J., et al., Twice-Yearly Depemokimab in Severe Asthma with an Eosinophilic Phenotype. N Engl J Med, 2024. 391(24): p. 2337–2349.

8. Wechsler, M.E., et al., Mepolizumab or Placebo for Eosinophilic Granulomatosis with Polyangiitis. N Engl J Med, 2017. 376(20): p. 1921–1932.

9. Mohammad, A.J., et al., Rituximab for the treatment of eosinophilic granulomatosis with polyangiitis (Churg-Strauss). Ann Rheum Dis, 2016. 75(2): p. 396–401.

10. James, L.K., B cells defined by immunoglobulin isotypes. Clin Exp Immunol, 2022. 210(3): p. 230–239.

11. King, H.W., et al., Integrated single-cell transcriptomics and epigenomics reveals strong germinal center-associated etiology of autoimmune risk loci. Sci Immunol, 2021. 6(64): p. eabh3768.

12. Kotagiri, P., et al., Disease-specific B cell clones are shared between patients with Crohn’s disease. Nat Commun, 2025. 16(1): p. 3689.

13. Galson, J.D., et al., Deep Sequencing of B Cell Receptor Repertoires From COVID-19 Patients Reveals Strong Convergent Immune Signatures. Front Immunol, 2020. 11: p. 605170.

14. Bashford-Rogers, R.J.M., et al., Analysis of the B cell receptor repertoire in six immune-mediated diseases. Nature, 2019. 574(7776): p. 122–126.

15. Piga, M.A., et al., Insights from comparison of serum IgG4 between T2-eosinophilic asthma and eosinophilic granulomatosis with polyangiitis/idiopathic hypereosinophilic syndrome. Clin Exp Rheumatol, 2025. 43(4): p. 710–717.

16. Sandeep, T., et al., Evaluation of serum immunoglobulin E levels in bronchial asthma. Lung India, 2010. 27(3): p. 138–40.

17. Oiwa, H., A. Yahata, and N. Nakamoto, Clinical value of biomarkers in relation to artery size in eosinophilic granulomatosis with polyangiitis: findings from an inception cohort at a Japanese City Hospital. Clin Rheumatol, 2025. 44(10): p. 4367–4373.

18. Ohm-Laursen, L., et al., B Cell Mobilization, Dissemination, Fine Tuning of Local Antigen Specificity and Isotype Selection in Asthma. Front Immunol, 2021. 12: p. 702074.

19. Asamori, T., et al., Molecular mimicry-driven autoimmunity in chronic rhinosinusitis with nasal polyps. J Allergy Clin Immunol, 2025. 155(5): p. 1521–1535.

20. Takatsu, K., Interleukin 5 and B cell differentiation. Cytokine Growth Factor Rev, 1998. 9(1): p. 25–35.

21. Buchheit, K.M., et al., *IL-5R*α *marks nasal polyp IgG4- and IgE-expressing cells in aspirin-exacerbated respiratory disease*. J Allergy Clin Immunol, 2020. 145(6): p. 1574–1584.

22. Loftus, B.G., et al., IgG subclass deficiency in asthma. Arch Dis Child, 1988. 63(12): p. 1434–7.

23. Singer, A., et al., Utility of immunology, microbiology, and helminth investigations in clinical assessment of severe asthma. J Asthma, 2022. 59(3): p. 541–551.

24. Klaustermeyer, W.B., et al., IgG subclass deficiency associated with corticosteroids in obstructive lung disease. Chest, 1992. 102(4): p. 1137–42.

25. Campbell, C.D., M. Gleeson, and I. Sulaiman, The role of the respiratory microbiome in asthma. Front Allergy, 2023. 4: p. 1120999.

26. Vidarsson, G., G. Dekkers, and T. Rispens, IgG subclasses and allotypes: from structure to effector functions. Front Immunol, 2014. 5: p. 520.

27. Zhang, Y., et al., Isolated IgG2 deficiency is an independent risk factor for exacerbations in bronchiectasis. QJM, 2022. 115(5): p. 292–297.

